# Symbiotic bacteria support calcium carbonate precipitation in the Gulf toadfish (*Opsanus beta*)

**DOI:** 10.1101/2025.10.07.681008

**Authors:** Anthony M. Bonacolta, Tristan Kravitz, Rocío Mozo, Lydia J. Baker, Rachael M. Heuer, Martin Grosell, Javier del Campo

## Abstract

Marine fish play a significant yet understudied role in the oceanic carbon cycle through the production of magnesium rich calcium carbonate (CaCO_3_) precipitates known as ichthyocarbonates. These deposits form in the gut of marine teleost fish in response to salinity, serving as part of their osmoregulation strategy. Through this, marine fish may contribute as much as 9.04 Pg of CaCO_3_ per year in global new carbonate production, being equivalent to or potentially higher than the production by coccolithophores and pelagic foraminifera. Despite their ecological relevance, the biological mechanisms driving ichthyocarbonate precipitation remain to be fully resolved. Intriguingly, bacteria are consistently found in intimate association with ichthyocarbonate precipitates. Given the widespread capacity of prokaryotes to mediate CaCO precipitation, this association points to a previously unexplored microbial contribution to the process. To investigate the potential role of bacteria in ichthyocarbonate production, we subjected Gulf toadfish (*Opsanus beta*) to a salinity challenge common to their native range and known to elicit elevated CaCO_3_ precipitation. To assess the respective contributions of the host and its microbiota to ichthyocarbonate formation in the gut, we characterized the microbiome across the toadfish gut and performed meta-transcriptomic analysis. Across the toadfish gut, we identify a high abundance of vibrios associated with ichthyocarbonates with the metabolic potential for CaCO_3_ precipitation. Specifically, we observe the expression of the transcriptional activator of urease (*ureR*) by *Photobacterium damselae* subsp*. damselae,* which can induce the precipitation of CaCO_3_ via the production of bicarbonate. We demonstrate that CaCO precipitation in marine fish may not solely be a host-driven process, but potentially the result of a functional symbiosis with gut-associated *Vibrio* bacteria. We hypothesize that just as photosymbionts enable corals to build reefs, fish hosts along with their microbial partners may synergistically contribute to oceanic carbonate production. This discovery, if confirmed, expands the role of symbiosis in marine biomineralization and underscores its broader influence on global biogeochemical cycles.

## Introduction

The marine environment is central to the global carbon cycle as the ocean is the largest reservoir of CO_2,_ which can be readily exchanged with the atmosphere, thus having a significant role in determining atmospheric CO_2_ concentrations over decadal to millennial timescales (1). Two mechanisms, the biological carbon pump and the carbonate pump, are integral to this process in the ocean. The biological carbon pump sequesters atmospheric CO_2_ in the form of photosynthetically fixed carbon, whereas the carbonate pump transfers CO_2_ back into the atmosphere via marine biogenic calcification. Marine fish play an important role not only in the biological carbon pump through prey consumption, fecal pellet production, and deadfall, but also in the carbonate pump through the production of calcium carbonate (CaCO_3_) precipitates (ichthyocarbonates), which form in their intestines from metabolic, endogenous CO_2_ derived from their diet (2,3). Through this process, marine fish are one of the dominant carbonate producers in the ocean, providing an estimated 0.33 to 9.03 Pg CaCO_3_ per year in global new carbonate production (3). This makes their contribution equivalent to or higher than more well-known sources such as coccolithophores and pelagic foraminifera (3–5). In contrast to those sources, marine fish carbonate excretion is anticipated to play an increasingly critical role in the inorganic carbon cycle under the developing conditions of global climate change as they increase ichthyocarbonate production in response to both rising temperatures and higher CO_2_ levels (6–9).

As mentioned above, marine fish produce ichthyocarbonates as part of their osmoregulatory strategy. Briefly, to maintain osmotic homeostasis while in a concentrated external environment, marine teleost fish ingest seawater to replace osmotic and urinary fluid loss (10). This seawater is first desalinated by the water-impermeable esophagus (11), followed by the anterior intestine, where NaCl uptake drives water absorption (12). This NaCl is later excreted across the gill epithelium (10). Bicarbonate (HCO_3_ ^−^) transported into the intestinal lumen facilitates water absorption (13). In the intestinal fluids CaCO_3_ is precipitated, removing Ca^2+^, Mg^2+^, and HCO_3_ ^−^ ions to lower the osmotic pressure. This precipitated CaCO_3_ then gets excreted into the marine environment. Several fish enzymes have been identified that play a role in this calcification process, including carbonic anhydrases, guanylin peptides, sodium-bicarbonate co-transporters, Na-H exchangers, V-type H-ATPases (14–16), but the CaCO_3_ biomineralization pathway remains incomplete within the fish genome. The role of microbes, specifically bacteria, in this calcification process has gone unexamined since the initial observation of bacteria being intimately associated with ichthyocarbonates three decades ago (17). Relatedly, CaCO_3_ precipitation has been extensively documented as a ubiquitous process among various bacterial groups in a diverse array of environments (18).

Symbiotic microbes can contribute to host physiology and its ability to adapt to certain environmental stressors (19), even co-evolving with their hosts over long periods (20). While microbiome research has gained traction in the last two decades, only recently have fish microbiomes gotten the same amount of attention, despite representing the greatest species diversity among vertebrates (21). Fish gut microbiomes are distinct from those of the water column and can vary significantly from fish to fish (22). Proteobacteria and Firmicutes are the dominant taxa of most fish gut microbiota, but environmental factors such as salinity play a major role in the microbiota composition (23,24). In marine fish, Proteobacteria are enriched compared to freshwater fish gut microbiomes. At the family level, *Moraxellaceae*, *Vibrionaceae*, *Enterobacteriaceae*, and *Alcaligenaceae* bacteria are significantly more common in marine fish compared to freshwater fish (23). Vibrios (family: *Vibrionaceae*), specifically, have been shown to dominate the gut microbiomes of marine fish species and also increase in abundance across a salinity gradient (25,26), indicating a likely physiological link between saltwater fish guts and these bacteria.

The Gulf toadfish, *Opsanus beta*, is an established marine teleost fish lab model, and the calcification process within their intestinal lumen has been extensively analyzed (13–15), making it the ideal subject for this investigation. We aim first to characterize the gut microbiome of the Gulf toadfish and then determine any possible role for the gut microbiota in the precipitation of ichthyocarbonates in this species. Using 16S rRNA gene metabarcoding, we profiled the microbiome of nine *O. beta* individuals across a gradient of salinity challenges (9 [brackish], 35 [marine], and 60 [hypersaline] parts per thousand [ppt]) designed to induce precipitate formation at the higher salinities, while also representing possible salinity levels this species may be exposed to in its natural habitat (27,28). Given that different microenvironments of the intestine serve distinct roles in ichthyocarbonate production (16), we split our sampling across four specific portions of the toadfish gut: the anterior intestine, posterior intestine, intestinal fluid, and ichthyocarbonates (when present; Figure 1A). This was followed by transcriptomic sequencing targeting bacteria and host reads from select samples to understand the metabolic contributions of each to CaCO_3_ precipitation. Phylogenetic analysis of genes involved in calcium carbonate precipitation confirmed the origin of essential genes detected in the transcriptomic data.

**Figure 1.**
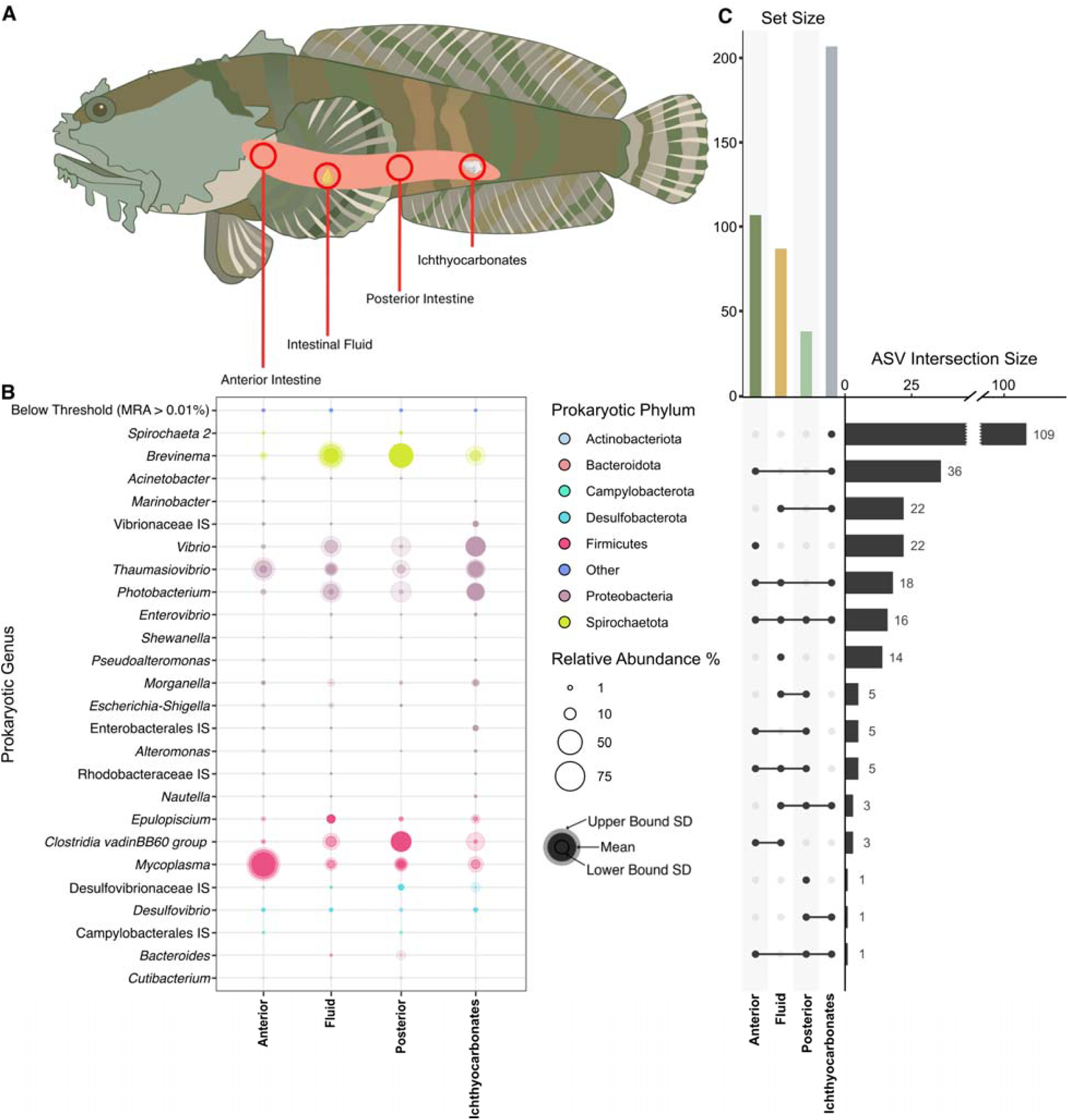
Prokaryotic community of the gulf toadfish (*O. beta*). **(A)** Diagram of sampled regions of the *O. beta* intestinal tract, including the anterior and posterior intestines, intestinal fluid, and ichthyocarbonates, when observed. Toadfish art adapted from Integration and Application Network (ian.umces.edu/media-library). **(B)** Bubble plot of prokaryotic genera across sampled regions. Only genera with a mean relative abundance (MRA) above 0.01% ar shown. Genera are colored by phylum, with upper bound and lower standard deviations indicated by bubble shading. IS – *incertae sedis*. (**C)** Upset plot depicting unique and shared ASVs across sampled regions of the Gulf toadfish gut. ASV intersection size represents ASVs common to each dot-connected grouping. Set size represents the total ASVs unique to each area.

## Results

1950 amplicon sequence variants (ASVs) were recovered across all samples. After filtering to just the prokaryotic ASVs present in the top 99% of at least one sample and with total counts above 150 reads, we were left with 436 prevalent ASVs for downstream analysis. Shannon- Weiner alpha diversity indices showed the prokaryotic community of each sampled intestinal region and ichthyocarbonates of *O. beta* to be significantly distinct from that of the water column (Supplementary Figure 1A). Aitchison Distance beta diversity showed no significant dissimilarity between sampled *O. beta* intestinal regions, but each region was distinct from the water column (Tukey-HSD; Supplementary Figure 1B, Supplementary Table 2).

In terms of microbiome composition, *Mycoplasma* spp. and *Thaumasiovibrio* spp. were highly abundant in anterior intestine samples across all salinity treatments (Figure 1B, Supplementary Figure 2). Intestinal fluid samples were dominated by Proteobacteria (*Vibrio* spp., *Thaumasiovibrio* spp., and *Photobacterium* spp.), Firmicutes (*Mycoplasma* spp., *Clostridia* spp., and *Epulopiscium* spp.), and spirochetes from the *Brevinema* genus (Figure 1B). The posterior intestine of *O. beta* exhibited a similar composition to the intestinal fluid; however, *Brevinema* spirochetes were more abundant in the higher salinity treatments (Supplementary Figure 2). As expected, ichthyocarbonates were only recovered in the higher salinity treatments and showed the most unique ASVs (Figure 1C). These precipitates exhibited a consistent dominance of proteobacteria, specifically *Photobacterium* at 35 ppt and *Vibrio* at 60 ppt (Figure 1B, Supplementary Figure 2). Only 16 ASVs were found across all *O. beta* samples (Figure 1C), with up to half of them belonging to Proteobacteria, specifically Gammaproteobacteria within the order Enterobacterales. Within this group, members of the *Vibrionaceae* family were highly prevalent, including representatives from the genera *Vibrio*, *Photobacterium*, and *Thaumasiovibrio*. Additionally, other Gammaproteobacteria such as *Alteromonas, Morganella* and *Shewanella* were also present in all samples. Although less abundant, additional shared phyla included Firmicutes (*Mycoplasma* spp.*, Epulopiscium* spp., and *Clostridia* spp.), Desulfobacterota (family *Desulfovibrionaceae*), Spirochaetota (*Brevinema* spp.), and Actinobacteriota (*Cutibacterium* spp.).

Analysis of compositions of microbiomes with bias correction (ANCOM-BC) confirmed that *Brevinema* sp. and *Vibrio* sp. were significantly abundant in ichthyocarbonate samples in comparison to the sampled parts of *O. beta* (padj = 1.69e-08 & 5.13e-06, respectively; Figure 2A). *Vibrio* sp., specifically, showed nearly ∼7.5 log fold change higher abundance in ichthyocarbonates (Figure 2A). Enterobacterales ASVs (predominantly from vibrios and *Photobacterium* spp.) dominate across all ichthyocarbonate samples, with ASV 5 (*Vibrio* sp.) making up nearly half the community at 60 ppt (Figure 2B). At 35 ppt, ASV 3 (*Thaumasiovibrio* sp.) and ASV 2 (*Photobacterium* sp.) dominated ichthyocarbonate samples (Figure 2B). The most abundant vibrios were placed onto a backbone reference tree of full-length Vibrionaceae 16S rRNA genes using RAxML’s evolutionary placement algorithm to confirm and further investigate their phylogenetic identity. ASV 5 was found to be most closely related to *Vibrio ponticus* and *Vibrio rhodolitus*. ASV 2 was nearly identical to *Photobacterium damselae*. ASV 3 and some other *O. beta* ASVs formed a distinct clade with *Vibrio stylophorae* (Figure 2C).

**Figure 2.**
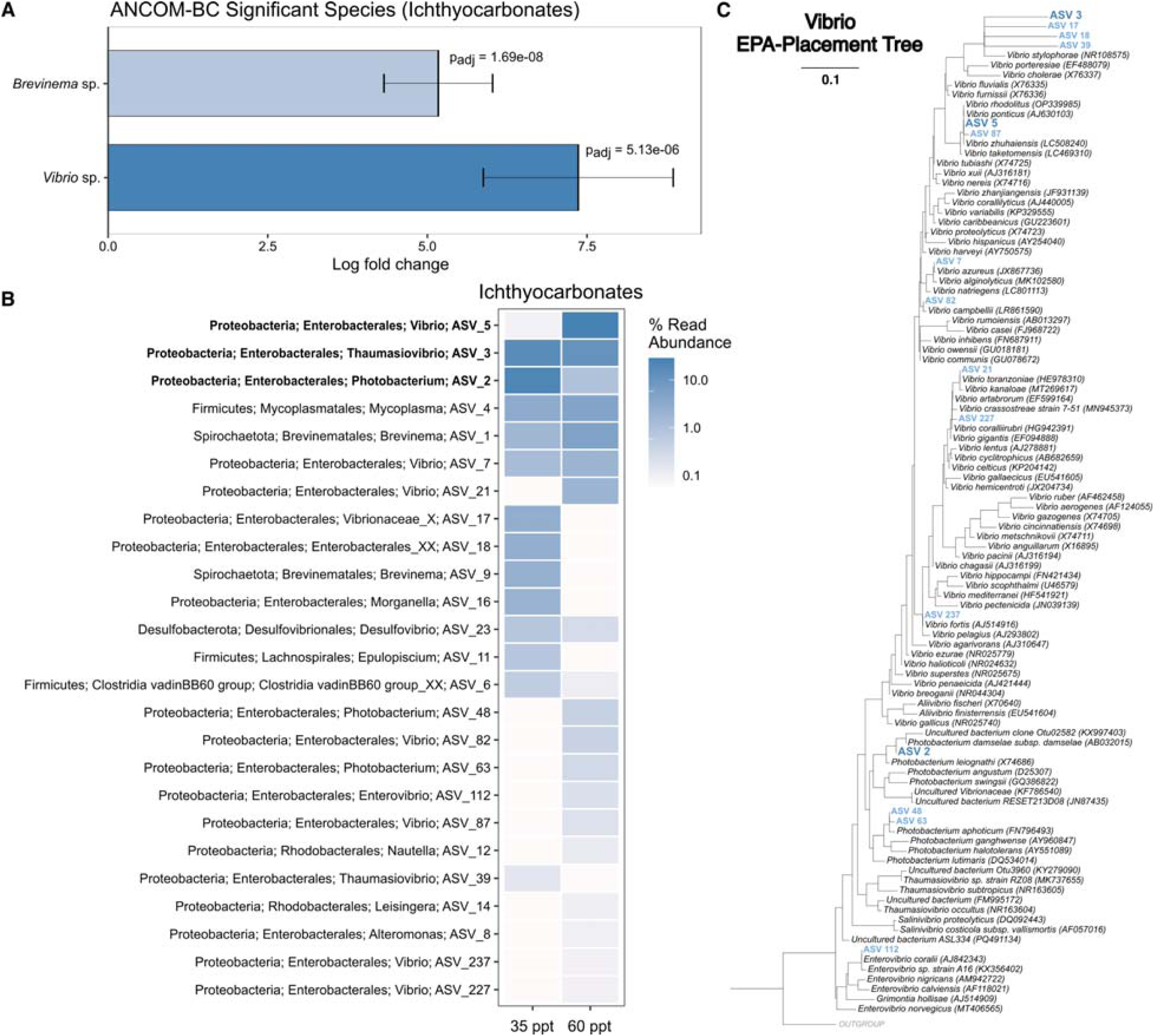
Abundant Vibrionaceae within the gulf toadfish (*O. beta*). **(A)** Analysis of compositions of microbiomes with bias correction (ANCOM-BC). Significant species from *O. beta* ichthyocarbonate samples vs. other sample regions. **(B)** Ampvis2 heatmap of the most abundant ASVs from *O. beta* ichthyocarbonates. The top three ASVs are bolded. (**C)** 16S rRNA gene phylogenetic tree of Vibrionaceae with ASVs from panel B placed using RAxML’ evolutionary-placement algorithm (EPA). Dark blue, bolded ASVs represent the 3 bolded ASV from panel B. Other Vibrionaceae ASVs from panel B are shown in lighter blue.

Genes hypothesized to be involved in CaCO_3_ precipitation across fish and bacteria were compiled following a literature search (Table 1). Assembled proteomes from RNA libraries of select samples were searched for the absence or presence of these genes. Phylogenetic analysis confirmed whether certain genes originated from *O. beta* or bacteria. All *O. beta* genes previously identified as important to the precipitation process were confirmed to be present in the host transcriptome (Table 1; see review by *Grosell and Oehlert 2023* (16)). Only *carbonic anhydrase* and *etfB* were found to be expressed by both the microbial community (not vibrios) and *O. beta*. Notably, we found that *fadR*, *fadB*, *ureR*, *napA*, and *napB* were expressed exclusively by the microbial community, specifically by *Vibrio* sp. or *Photobacterium* sp. associated with ichthyocarbonates (Table 1). Phylogenetic analyses confirmed that these genes were not expressed by *O. beta*. The *ureR* gene recovered from ichthyocarbonate samples branched with and was nearly identical to *ureR* expressed by *Photobacterium damselae* (Figure 3A). Despite not being able to recover urease accessory genes from the transcriptomic data, Phylogenetic Investigation of Communities by Reconstruction of Unobserved States (PICRUSt2) analysis of the microbial community confirmed a high genomic potential of these genes in ichthyocarbonates.

**Figure 3.**
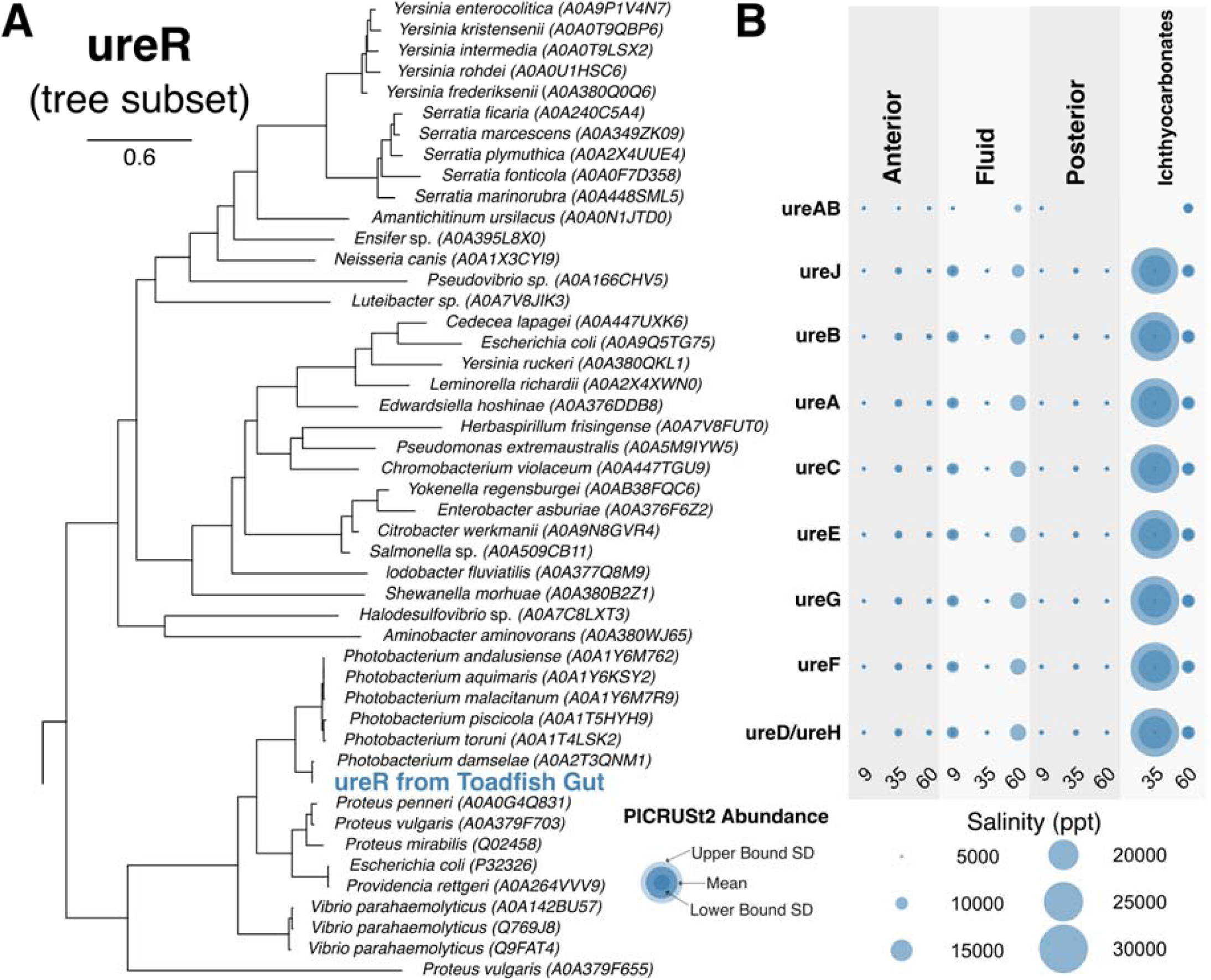
Urease gene expression and metabolic potential of O. beta intestinal microbiota. **(A)** Phylogenetic tree of ureR (transcriptional activator of the urease operon), including the *Photobacterium damselae*-related ureR detected from the gut of *O. beta*. Accession numbers are shown in parentheses. **(B)** Functional abundance of urease accessory genes across the sampled regions of *O. beta* in each salinity treatment according to a PICRUSt2 (Phylogenetic Investigation of Communities by Reconstruction of Unobserved States) analysis. Upper bound and lower bound standard deviation is indicated by bubble shading.

**Table 1.** Genes involved in CaCO_3_ precipitation detected within our data.

| *Gene* | *Toadfish* | *Bacteria* | *Vibrio* | *Reference* | *Notes* |
| --- | --- | --- | --- | --- | --- |
| *fadR* (*ysiA*) / *Fatty acid metabolism regulator protein* | - | + | +<br>*Photobacterium* | (29) | Identified as essential for precipitation process in *Bacillus subtilis* |
| *fadB* (*ysiB*) / *Fatty acid oxidation complex subunit alpha* | - | + | +<br>*Vibrios and Photobacterium* | (29) | Identified as essential for precipitation process in *Bacillus subtilis* |
| *Electron transfer flavoprotein subunit beta* (*etfB*) | + | + | - | (29) | Identified as essential for precipitation process in *Bacillus subtilis* |
| *Carbonic anhydrase* (*CA*) | + | + | - | (30) | Catalyzes the reversible hydration of carbon dioxide: CO<sub>2</sub> + H<sub>2</sub>O ↔ HCO<sub>3</sub><sup>-</sup> + H<sup>+</sup>, plays a role in increasing bicarbonate concentration |
| *Urease* (*Ure*) | - | + | +<br>*Photobacterium damsela* | (31) | Releases ammonia and increases urinary pH via urea hydrolysis, resulting in the precipitation of calcium and magnesium compounds into urinary stones |
| *Periplasmic nitrate reductase* (*napAB*) | - | + | +<br>*Vibrios and Photobacterium* | (32) | Nitrate reduction and ammonification of amino-acids are prone to induce the precipitation of CaCO<sub>3</sub> |
| *Respiratory nitrate reductase* (*narGHI*) | - | + | - | (32) | Nitrate reduction and ammonification of amino-acids are prone to induce the precipitation of CaCO<sub>3</sub> |
| *electrogenic* (HCO<sub>3</sub><sup>-</sup>:Cl<sup>-</sup>) *anion exchanger* (*SLC26a6*) | + | - | - | (33,34) | Intestinal bicarbonate excretion occurs along the entire length of the fish intestine via this protein, resulting in high luminal bicarbonate concentrations |
| *Na<sup>+</sup>:HCO<sub>3</sub><sup>-</sup> cotransporter* (*NBC1 / SLC4A7*) | + | - | - | (35,36) | Transports bicarbonate through the fish epithelial cells of the intestine |
| *Na<sup>+</sup>:K<sup>+</sup>-ATPase* / *Sodium potassium pump* | + | - | - | (36,37) | Fuels bicarbonate transporters in the fish gut |
| *Na<sup>+</sup>:K<sup>+</sup>:2Cl<sup>-</sup> cotransporter* (*NKCC/ SLC12A1*) | + | - | - | (38–41) | Mediates guanylin cyclase-initiated secretion of chloride ions in the distal region of the fish intestinal tract |
| *soluble adenyl cyclase* (*sAC*) | + | - | - | (42) | Stimulates NKCC2 under conditions of high luminal bicarbonate concentrations |
| *Guanylin peptides* (*GN*) / *Guanylin cyclase-C* (*CG-C*) | + | - | - | (38–41) | Found throughout the fish intestine and facilitates excretion of ichthyocarbonates |
| *cystic fibrosis transmembrane regulator Cl<sup>-</sup> channel* (*CFTR*) | + | - | - | (38–41) | Mediates guanylin cyclase-initiated secretion of chloride ions in the distal region of the fish intestinal tract |

## Discussion

This is the first study in 30 years to investigate the potential contribution of microbes to ichthyocarbonate production. We identified both the microbes and prokaryotic gene expression capable of supporting CaCO_3_ precipitation in the gut of *O. beta*. The prokaryotic microbiome of *O. beta* was stable across sampled regions of the intestine and distinct from that of the tank water (Figure 1B & Supplementary Figure 2). Consistent with previous literature analyzing the gut composition of marine fish, we saw Proteobacteria (especially vibrios) and Firmicutes (*Mycoplasma* spp.) as significant members of the intestinal community (23,24) (Figure 1B). *Mycoplasma* spp. have previously been observed in clusters on the intestinal mucosa of fish and appear adapted to the nutrient-rich environment of the gut (43). *Brevinema* spirochetes, which were most common in fluid and posterior intestine samples (Figure 1B), have also been noted as common inhabitants of fish guts before and are presumed to have a similar nutrient scavenging role (43,44). Importantly, we note the high abundance of various Vibrionaceae within the gut of *O. beta,* including *Vibrio* spp., *Photobacterium* spp., and *Thaumasiovibrio* spp. (Figure 1B, Figure 2A-C). The surface of the ichthyocarbonates, which form in higher salinities as part of the toadfish’s osmoregulatory process, is dominated by vibrios, specifically ASVs related to *V. ponticus, V. rhodolitus*, *P. damselae*, and *V. stylophorae* (Figure 2A-C). *V. ponticus* and *V. rhodolitus* are members of the Ponticus vibrio clade (45). Members of this clade have been isolated from diseased fish, crustaceans, and bivalves (46), however, we did not note any signs of illness within our sampled fish. *P. damselae* has also been reported as a pathogen of a variety of marine animals and even in humans (47); however, it has also been collected from marine fish not showing any signs of disease (48), like the *O. beta* used in this study. *V. stylophorae* was first isolated from a reef-building coral in Taiwan (49). The unique clustering of our fish-associated *Vibrio* ASVs as sister to *V. stylophorae* suggests a possible fish host-specific divergence in this lineage.

Along with the high abundance of vibrios on ichthyocarbonates, we also identify vibrio-specific gene expression within these samples that would aid in the precipitation of calcium carbonate. Two of these genes, *fadR* and *fadB,* are synonymous to the *ysiA* and *ysiB* genes in *Bacillus subtilis*, which have been shown to induce calcium carbonate biomineralization (29). As these genes are involved in fatty acid metabolism, it is thought that an intermediate in this process likely contributes to the biomineralization process. Likewise, *napA* and *napB,* which encode periplasmic nitrate reductases, were also found to be expressed by vibrios within the gut of the gulf toadfish (Table 1). Furthermore, *narG* and *narH*, which encode respiratory nitrate reductases, were expressed by non-Vibrio bacteria within the gut of *O. beta* as well (Table 1). As nitrate reduction coupled with an organic carbon source generates CO_2,_ which equilibrates with water to form bicarbonate ions, the expression of these genes by the intestinal microbiota of *O. beta* is capable of promoting CaCO_3_ precipitation (32). More directly, the genomic potential of urease accessory proteins combined with the observed expression of *ureR* (urease operon transcriptional activator) by vibrios (Figure 3), specifically *P. damselae,* which we also found as highly abundant on ichthyocarbonates, supports a microbial contribution to ichthyocarbonate production. Urease is the enzyme that catalyzes the hydrolysis of urea into ammonia and carbamic acid. Carbamic acid spontaneously decomposes into additional ammonia and carbonic acid (50). Carbonic acid dissociates to form bicarbonate, which likely reacts with Ca^2+^ ions within the fish gut to precipitate CaCO_3_ (Figure 3B). The ability to hydrolyze urea is a key characteristic of *P. damselae* subsp*. damselae,* which discriminates it from *P. damselae* subsp. *piscicida* (51), further discerning ASV 2 as most likely *P. damselae* subsp*. damselae*. This *P. damselae* pathway, combined with the expression of *carbonic anhydrases* (which catalyzes the hydration of carbon dioxide, increasing bicarbonate concentration (30)) by the toadfish host, further induces ichthyocarbonate production. Vibrios have been hypothesized to play an active role in CaCO_3_ precipitation in their natural habitats (52), and here we show for the first time the potential for this within the fish gut microenvironment.

Taken together, our data support the role of intestinal microbiota, specifically vibrios, in the precipitation of CaCO_3_ in the guts of marine fish. The abundance of *P. damselae* subsp. *damselae* on ichthyocarbonates and their expression of *ureR* suggest a partnership with the fish host, whereby microbial production of bicarbonate works alongside host enzymatic activity to drive CaCO_3_ precipitation. We also report several other vibrio-encoded genes involved in fatty acid metabolism (*fadR* & *fadB*) and nitrate reduction (*napA* & *napB*), which also contribute to biomineralization conditions in fish intestines. The interplay between host enzymes and gut-associated vibrios creates the necessary conditions for CaCO_3_ precipitation, unveiling a novel symbiotic pathway in marine biomineralization. Based on these findings, we hypothesize that fish gut bacteria may function as significant microbial contributors to global carbonate cycling alongside coccolithophores, foraminifera, and coral symbionts. Future research should confirm these findings by directly measuring the contribution of vibrio-derived bicarbonate to ichthyocarbonate composition and clarify the prevalence and specificity of this partnership across fish species.

## Materials and Methods

### Sample Collection

Gulf toadfish were obtained as bycatch from local shrimp fishermen in Biscayne Bay, FL, and transported to laboratory holding tanks following brief (5 min) freshwater (FW) baths and malachite green treatment to treat for any ectoparasites (36). Holding tanks (62 L) were continuously provided with flow-through, sand-filtered seawater (SW) from Biscayne Bay (32– 35 ppt salinity, 25–32°C) and constant aeration. Gulf toadfish were fed to satiation twice weekly with meals consisting of chopped squid and shrimp. After approximately one month in holding tanks, Gulf toadfish were transferred to experimental tanks (62 L) containing either sand-filtered Biscayne Bay SW, 60 ppt or 9 ppt water and were allowed to acclimate for 1 week. 60 ppt water was produced by adding Instant Ocean sea salt to seawater while 9 ppt water was produced by diluting seawater with deionized water. Common for all three salinity treatments was renewal of the water every second day and constant aeration.

Samples were obtained from the intestinal lumen and from epithelial scrapings of individual toadfish exposed to different salinity treatments (n = 3 fish per treatment, mass = 95.5 ± 3.9 g). Fish were kept at three different salinity treatments because salinity has been observed to impact carbonate precipitation (13). At 9 ppt, no ichthyocarbonates were found to have formed. Whereas the fish kept at 35 ppt and 60 ppt salinity, corresponding to normal seawater and hyper-salinity, had ample ichthyocarbonate production. *O. beta* sampling occurred at four locations: the anterior intestine, posterior intestine, intestinal fluid, and ichthyocarbonates (Figure 1A). Water samples were also collected from each respective salinity treatment. Samples were extracted using the Qiagen All Prep DNA/RNA Minikit (Qiagen, Hilden, Germany).

### 16S rRNA Gene Metabarcoding

The V4 region of the 16S rRNA gene was amplified according to the Earth Microbiome protocol using 515F (5’ – GTGYCAGCMGCCGCGGTAA – 3’) and 806R (5’ – GGACTACNVGGGTWTCTAAT – 3’) primers from Apprill and Parada (53,54). PCR mixture for one sample contained 10.0 μl PCR master mix (2x), 0.5 μl each of the forward and reverse primers (diluted at 1:10), 1.0 μl template DNA, and reaction volumes made up to 25.0 μl with PCR-grade water (13.0 μl). The PCR program of the thermal cycler was set up as follows: an initial denaturation step at 94°C for 3 minutes, subsequent 35 cycles of denaturation at 94°C for 45 seconds, 35 cycles of annealing at 50°C for 60 seconds, and 35 cycles of elongation at 72°C for 90 seconds, all of them followed by a final elongation at 72°C for 10 minutes. Amplification was checked by gel electrophoresis. Electrophoresis gel was made with 1g of ultrapure agarose dissolved in 100 ml of 1X Tris-acetate-EDTA buffer (TAE) and stained with 5 μl of SYBR safe DNA gel stain. The gel was run for about 20 minutes at 100 V at room temperature, then visualized under a blue light transilluminator. Amplified PCR products were cleaned using AMPure magnetic beads (Thermofisher). DNA concentrations from purified PCR samples were checked using a Qubit fluorometer (Thermofisher) before being sent to the Integrated Microbiome Resource facility at the Centre for Comparative Genomics and Evolutionary Bioinformatics at Dalhousie University for sequencing on an Illumina MiSeq using 300+300 bp paired end V3 chemistry. Samples that failed the first round of sequencing were further purified using AMPure magnetic beads and sent to Novogene for another round of sequencing on an Illumina MiSeq following the same library prep instructions.

### Microbiome Analysis

Primers and low-quality bases were removed from reads using Cutadapt v3.1 (55). The trimmed reads were then processed in R using DADA2 (56). Chimeras were removed using the ‘removeBimeraDenovo’ command with the “consensus” option. For the 16S rRNA gene amplicons, forward reads were truncated at 250 bp and reverse reads at 210 bp, corresponding to a general drop-off in read quality past these points of the sequences. Truncated reads were then denoised and merged into amplicon sequence variants (ASVs) before assigning taxonomy using SILVA v138 in DADA2 (57). Sequence tables, taxonomy tables, and metadata for the 16S rRNA gene amplicon datasets were uploaded into a Phyloseq object in R for further filtering and analysis (58). For the 16S rRNA gene amplicons, ASVs corresponding to Chloroplast and Mitochondria were removed. Furthermore, only ASVs present in the top 99% of at least one sample and with total counts above 150 reads were kept for further analysis. Bubble plots, alpha-diversity, and beta-diversity figures were constructed using ggplot2 and tidyverse packages (59). Statistical analyses were conducted using vegan (60). Analysis of similarities (ANOSIM) within vegan was used to test for differences in beta-diversity between groups using 999 permutations. Ampvis2 aided in distinguishing the most abundant and significant ASVs (61). Analysis of Compositions of Microbiomes with Bias Correction (ANCOM-BC) was used to find significantly enriched and differentially abundant taxa between treatment groups while adjusting for a false discovery rate (62). PICRUSt2 was run on the filtered ASV dataset to assess the metagenomic potential of the prokaryotic community (63).

### Vibrio Phylogenetics and Evolutionary Placement of ASVs

Phylogenetic analyses of the family *Vibrionaceae* were conducted to assess the evolutionary relationships between abundant ASVs from the ichthyocarbonates and closely related species. Due to the short length of the ASV sequences, these were placed onto a previously generated 16S rRNA maximum likelihood phylogenetic tree using the Evolutionary Placement Algorithm (EPA). To construct the 16S phylogenetic tree, we compiled a dataset comprising a total of 87 *Vibrionaceae* sequences and 3 *Shewanellaceae* sequences as an outgroup. These context sequences were downloaded from the NCBI database. The 90 sequences were aligned using MAFFT v7.505 (64), and the alignment was inspected in AliView v1.28 (65). For clean-up of the alignment, we employed trimAl v1.4.rev15 (66), which performed an automated alignment trimming. A maximum likelihood phylogenetic analysis was then performed using RAxML v8.2.12 (67) to construct the final *Vibrionaceae* phylogenetic tree. The most abundant *Vibrio* ASVs from the ichthyocarbonates were subsequently placed into this backbone reference tree using RAxML’s EPA. For this, the short ASV sequences were first aligned to the existing *Vibrionaceae* 16S rRNA gene alignment using MAFFT v7.505. The EPA analysis was then run to position the ASVs within the phylogeny. Finally, the resultant EPA tree was visualized and edited using the Interactive Tree of Life (iTOL) online platform.

### Host and Microbial Transcriptomic Analysis

Select samples were chosen for either host or prokaryotic community RNA sequencing through Novogene, depending on sample type and RNA quality (Supplementary Table 1). This resulted in 5 anterior gut samples for host mRNA sequencing via poly-A selection and 2 ichthyocabonate samples for prokaryotic mRNA sequencing via ribosomal depletion. Sequencing was conducted through Novogene on an Illumina NovaSeq targeting 6 GB of 150+150 PE reads per sample. Trim Galore! (68) was used to trim adaptors and poor-quality reads. Ribosomal RNAs were removed with sortmeRNA (69). Cleaned reads were then aligned to a *O. beta* genomic reference using STAR to generate gene counts (70,71). For the prokaryotic RNA samples, the unmapped reads were assembled using rnaSPAdes (72). ORFs were identified, and then protein prediction was performed using TransDecoder with BlastP and hmmscan with the uniport and pfam databases (73–75). Proteins were annotated using e-mapper (76). Detected urease proteins were confirmed through SGTs. Briefly, SGTs were generated by first generating a protein alignment using linsi (64). Alignments were trimmed with trimal and phylogenetic trees constructed using FastTree (66,77). Hmmsearch was also used to detect urease genes within the predicted proteomes.

## Data availability

The raw reads for the project have been deposited on NCBI SRA (BioProject: PRJNA1308130). Code used for transcriptomic analysis, single-gene trees, as well as ASV count tables, taxonomy tables, and nucleotide sequences can be found on GitHub at: https://github.com/delCampoLab/toadfish_microbiome.

## Acknowledgments

M.G. is a Maytag Professor of Ichthyology. The authors acknowledge Caleigh Weis for assistance with DNA clean-up for some microbiome samples. This work was supported by start-up funds from the University of Miami.

---

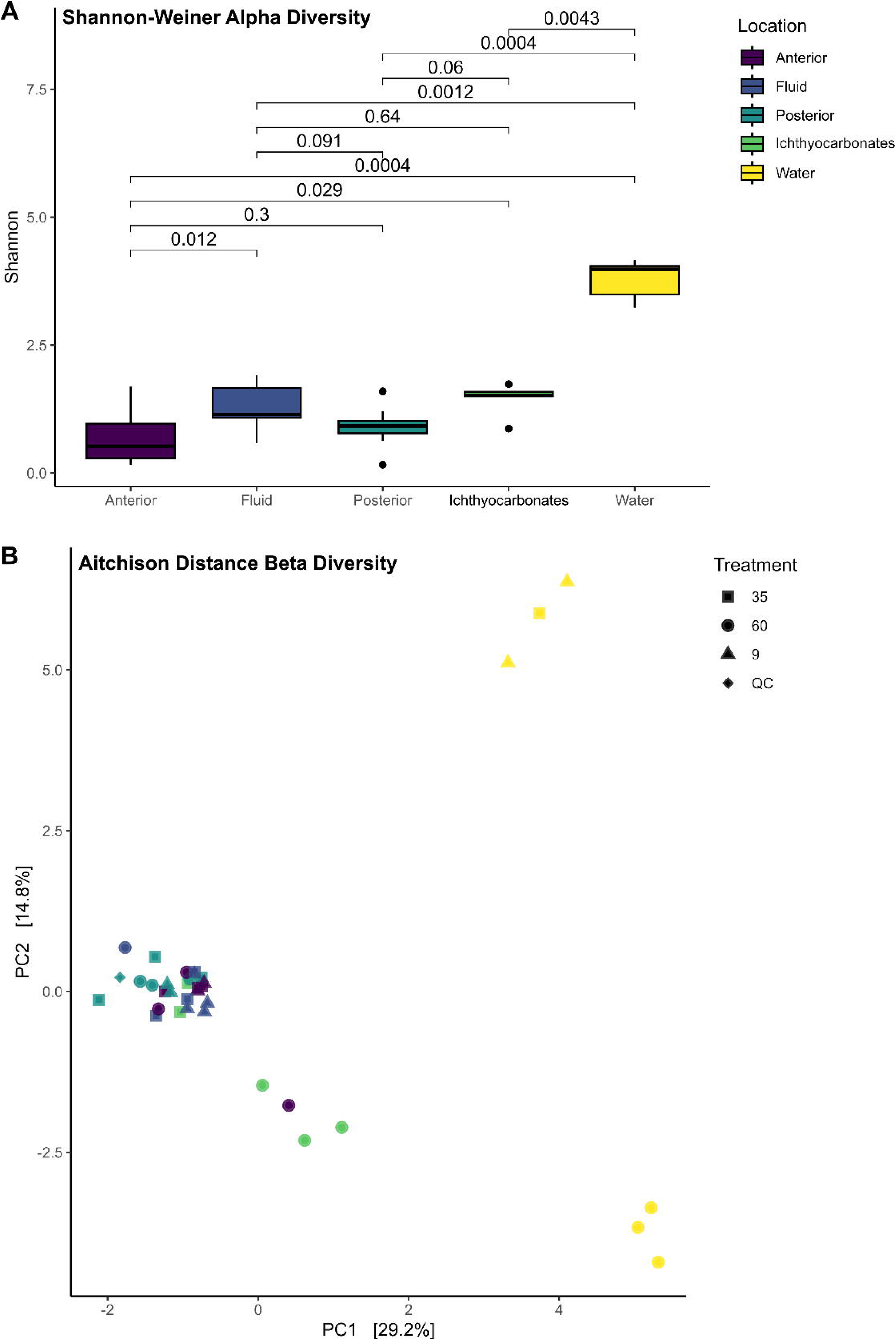

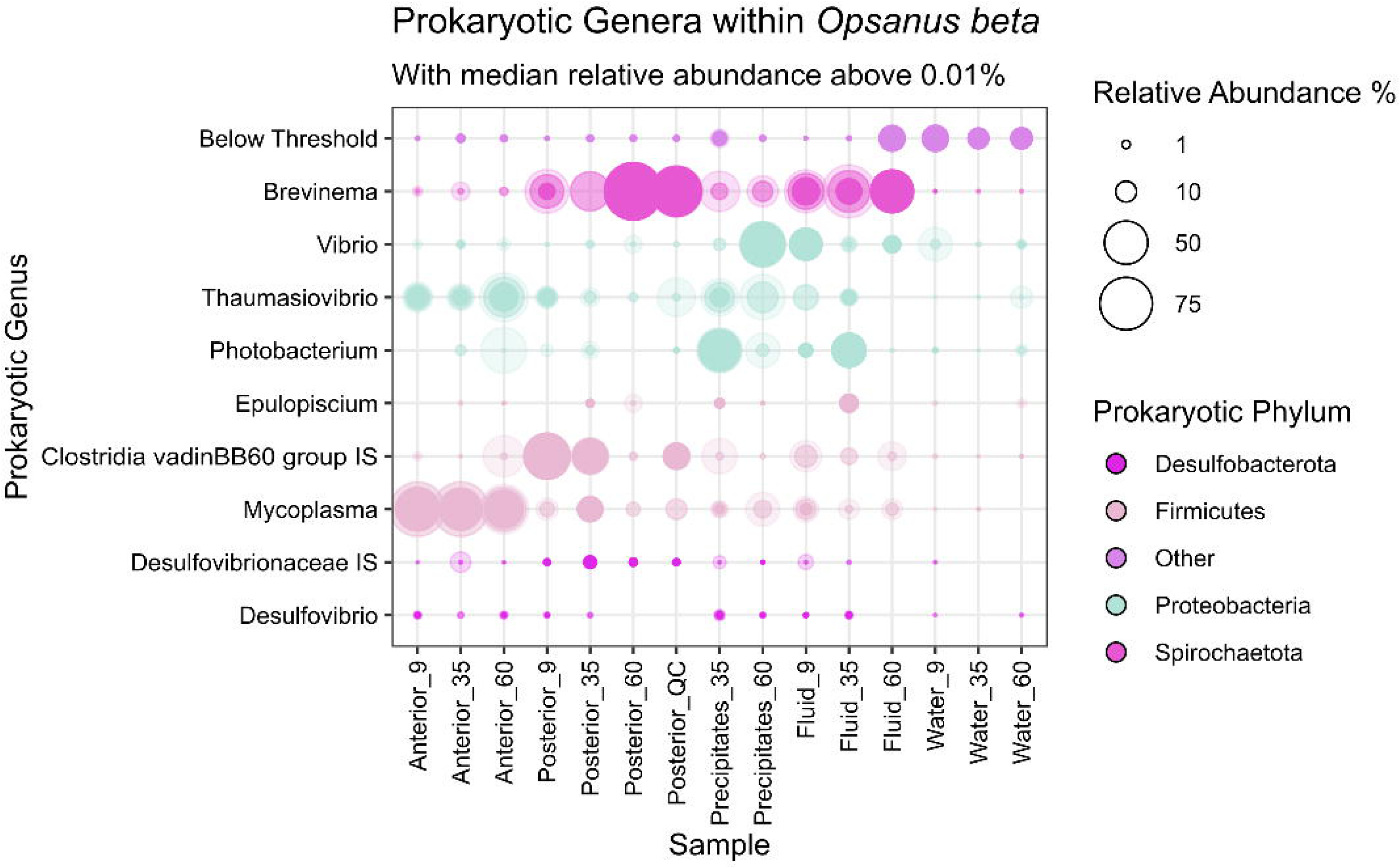

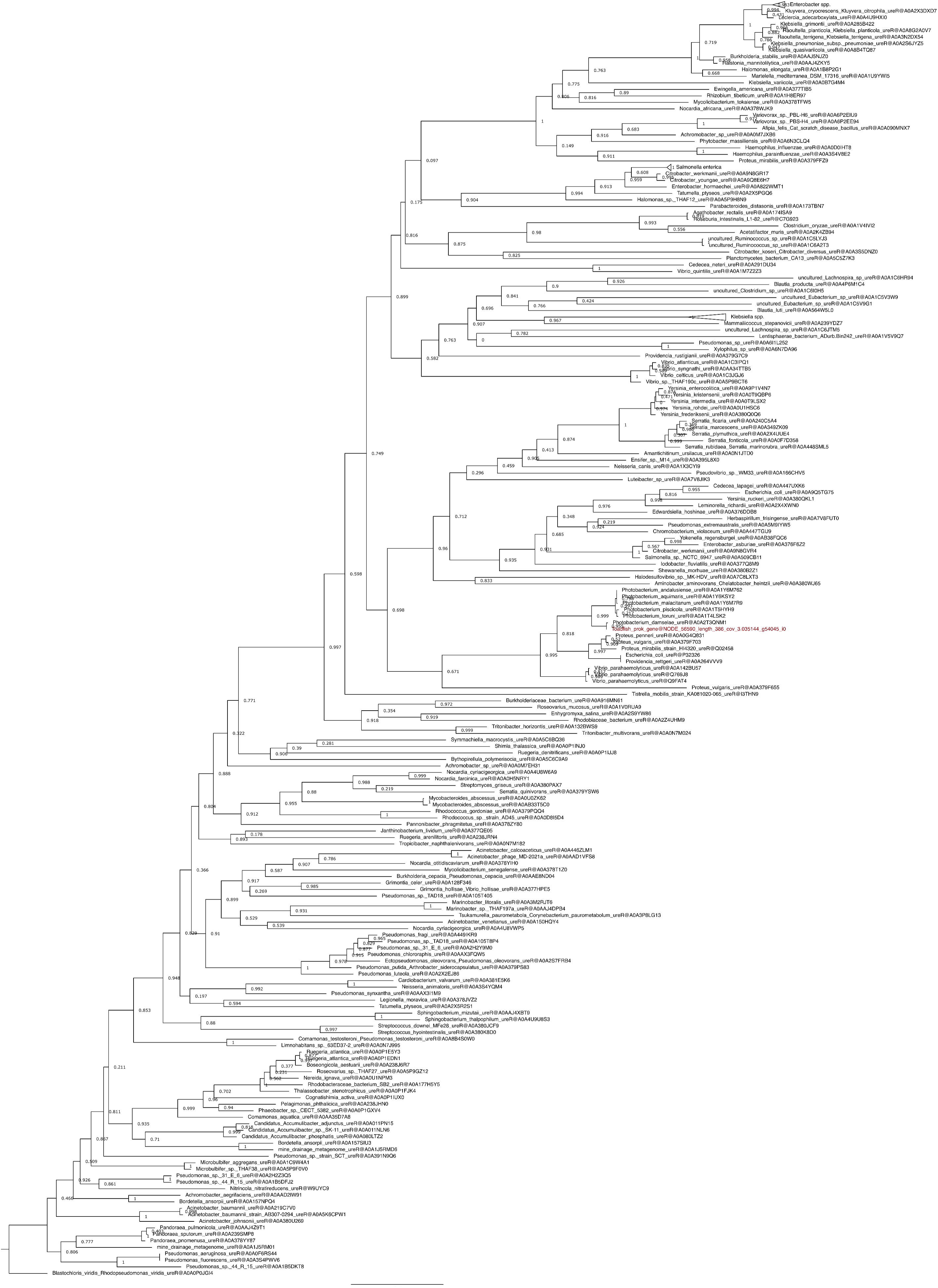

## References

1. DeVries T. The Ocean Carbon Cycle. Annu Rev Environ Resour. 2022 Oct 17;47(Volume 47, 2022):317–41.

2. Grosell M, Laliberte CN, Wood S, Jensen FB, Wood CM. Intestinal HCO3- secretion in marine teleost fish: Evidence for an apical rather than a basolateral Cl-/HCO3- exchanger. Fish Physiol Biochem. 2001;24(2):81–95.

3. Oehlert AM, Garza J, Nixon S, Frank L, Folkerts EJ, Stieglitz JD, et al. Implications of dietary carbon incorporation in fish carbonates for the global carbon cycle. Sci Total Environ. 2024 Mar 15;916:169895.

4. Milliman JD, Droxler AW. Neritic and pelagic carbonate sedimentation in the marine environment: ignorance is not bliss. Geol Rundsch. 1996 Sep 1;85(3):496–504.

5. Godrijan J, Drapeau DT, Balch WM. Osmotrophy of dissolved organic carbon by coccolithophores in darkness. New Phytol. 2022;233(2):781–94.

6. Grosell M. CO2 and calcification processes in fish. In: Grosell M, Munday PL, Farrell AP, Brauner CJ, editors. Fish Physiology [Internet]. Academic Press; 2019 [cited 2025 Mar 26]. p. 133–59. (Carbon Dioxide; vol. 37). Available from: https://www.sciencedirect.com/science/article/pii/S1546509819300020

7. Alves A, Gregório SF, Ruiz-Jarabo I, Fuentes J. Intestinal response to ocean acidification in the European sea bass (*Dicentrarchus labrax*). Comp Biochem Physiol A Mol Integr Physiol. 2020 Dec 1;250:110789.

8. Gregório SF, Ruiz-Jarabo I, Carvalho EM, Fuentes J. Increased intestinal carbonate precipitate abundance in the sea bream (Sparus aurata L.) in response to ocean acidification. PLOS ONE. 2019 Jun 21;14(6):e0218473.

9. Heuer RM, Munley KM, Narsinghani N, Wingar JA, Mackey T, Grosell M. Changes to Intestinal Transport Physiology and Carbonate Production at Various CO2 Levels in a Marine Teleost, the Gulf Toadfish (Opsanus beta). Physiol Biochem Zool PBZ. 2016;89(5):402–16.

10. Larsen EH, Deaton LE, Onken H, O’Donnell M, Grosell M, Dantzler WH, et al. Osmoregulation and Excretion. In: Comprehensive Physiology [Internet]. John Wiley & Sons, Ltd; 2014 [cited 2025 Mar 26]. p. 405–573. Available from: https://onlinelibrary.wiley.com/doi/abs/10.1002/cphy.c130004

11. Hirano T, Mayer-Gostan N. Eel esophagus as an osmoregulatory organ. Proc Natl Acad Sci. 1976 Apr;73(4):1348–50.

12. Grosell M, Taylor JR. Intestinal anion exchange in teleost water balance. Comp Biochem Physiol A Mol Integr Physiol. 2007 Sep 1;148(1):14–22.

13. Schauer KL, Reddam A, Xu EG, Wolfe LM, Grosell M. Interrogation of the Gulf toadfish intestinal proteome response to hypersalinity exposure provides insights into osmoregulatory mechanisms and regulation of carbonate mineral precipitation. Comp Biochem Physiol Part D Genomics Proteomics. 2018 Sep;27:66–76.

14. Schauer KL, LeMoine CMR, Pelin A, Corradi N, McDonald MD, Warren WC, et al. A proteinaceous organic matrix regulates carbonate mineral production in the marine teleost intestine. Sci Rep. 2016 Dec 23;6(1):34494.

15. Schauer KL, Christensen EAF, Grosell M. Comparison of the organic matrix found in intestinal CaCO3 precipitates produced by several marine teleost species. Comp Biochem Physiol A Mol Integr Physiol. 2018 Jul 1;221:15–23.

16. Grosell M, Oehlert AM. Staying Hydrated in Seawater. Physiology. 2023 Jul;38(4):178–88.

17. Walsh PJ, Blackwelder P, Gill KA, Danulat E, Mommsen TP. Carbonate deposits in marine fish intestines: A new source of biomineralization. Limnol Oceanogr. 1991;36(6):1227–32.

18. Perito B, Mastromei G. Molecular basis of bacterial calcium carbonate precipitation. Prog Mol Subcell Biol. 2011;52:113–39.

19. Voolstra CR, Ziegler M. Adapting with Microbial Help: Microbiome Flexibility Facilitates Rapid Responses to Environmental Change. BioEssays. 2020;42(7):2000004.

20. Ley RE, Lozupone CA, Hamady M, Knight R, Gordon JI. Worlds within worlds: evolution of the vertebrate gut microbiota. Nat Rev Microbiol. 2008 Oct;6(10):776–88.

21. Nelson JS, Grande TC, Wilson MVH. Fishes of the World. John Wiley & Sons; 2016. 1067 p.

22. Gallo BD, Farrell JM, Leydet BF. Fish Gut Microbiome: A Primer to an Emerging Discipline in the Fisheries Sciences. Fisheries. 2020 May 1;45(5):271–82.

23. Kim PS, Shin NR, Lee JB, Kim MS, Whon TW, Hyun DW, et al. Host habitat is the major determinant of the gut microbiome of fish. Microbiome. 2021 Jul 31;9(1):166.

24. Sullam KE, Essinger SD, Lozupone CA, O’connor MP, Rosen GL, Knight R, et al. Environmental and ecological factors that shape the gut bacterial communities of fish: a meta-analysis. Mol Ecol. 2012;21(13):3363–78.

25. Smriga S, Sandin SA, Azam F. Abundance, diversity, and activity of microbial assemblages associated with coral reef fish guts and feces. FEMS Microbiol Ecol. 2010 Jul 1;73(1):31–42.

26. Schmidt VT, Smith KF, Melvin DW, Amaral-Zettler LA. Community assembly of a euryhaline fish microbiome during salinity acclimation. Mol Ecol. 2015;24(10):2537–50.

27. Kelble CR, Johns EM, Nuttle WK, Lee TN, Smith RH, Ortner PB. Salinity patterns of Florida Bay. Estuar Coast Shelf Sci. 2007 Jan 1;71(1):318–34.

28. Lee TN, Johns E, Melo N, Smith RH, Ortner P, Smith D. On Florida Bay hypersalinity and water exchange. Bull Mar Sci. 2006 Sep 1;79(2):301–27.

29. Barabesi C, Galizzi A, Mastromei G, Rossi M, Tamburini E, Perito B. Bacillus subtilis Gene Cluster Involved in Calcium Carbonate Biomineralization. J Bacteriol. 2007 Jan;189(1):228–35.

30. Unzueta-Martínez A, Girguis PR. Taxonomic diversity and functional potential of microbial communities in oyster calcifying fluid. Appl Environ Microbiol. 2024 Dec 12;91(1):e01094–24.

31. Mobley HL, Island MD, Hausinger RP. Molecular biology of microbial ureases. Microbiol Rev. 1995 Sep;59(3):451–80.

32. Jain S, Fang C, Achal V. A critical review on microbial carbonate precipitation via denitrification process in building materials. Bioengineered. 12(1):7529–51.

33. Grosell M, Genz J, Taylor JR, Perry SF, Gilmour KM. The involvement of H+- ATPase and carbonic anhydrase in intestinal HCO3- secretion in seawater- acclimated rainbow trout. J Exp Biol. 2009 Jun;212(Pt 12):1940–8.

34. McDonald MD. The Renal Contribution to Salt and Water Balance. In: Fish Osmoregulation. CRC Press; 2007.

35. Chang MH, Plata C, Kurita Y, Kato A, Hirose S, Romero MF. Euryhaline pufferfish NBCe1 differs from nonmarine species NBCe1 physiology. Am J Physiol-Cell Physiol. 2012 Apr 15;302(8):C1083–95.

36. Taylor JR, Mager EM, Grosell M. Basolateral NBCe1 plays a rate-limiting role in transepithelial intestinal HCO3– secretion, contributing to marine fish osmoregulation. J Exp Biol. 2010 Feb 1;213(3):459–68.

37. Grosell M, Genz J. Ouabain-sensitive bicarbonate secretion and acid absorption by the marine teleost fish intestine play a role in osmoregulation. Am J Physiol-Regul Integr Comp Physiol. 2006 Oct;291(4):R1145–56.

38. Ruhr IM, Bodinier C, Mager EM, Esbaugh AJ, Williams C, Takei Y, et al. Guanylin peptides regulate electrolyte and fluid transport in the Gulf toadfish (Opsanus beta) posterior intestine. Am J Physiol-Regul Integr Comp Physiol. 2014 Nov;307(9):R1167–79.

39. Ruhr IM, Mager EM, Takei Y, Grosell M. The differential role of renoguanylin in osmoregulation and apical Cl−/HCO3− exchange activity in the posterior intestine of the Gulf toadfish (Opsanus beta). Am J Physiol-Regul Integr Comp Physiol. 2015 Aug 15;309(4):R399–409.

40. Ruhr IM, Schauer KL, Takei Y, Grosell M. Renoguanylin stimulates apical CFTR translocation and decreases HCO3− secretion through PKA activity in the Gulf toadfish (Opsanus beta). J Exp Biol. 2018 Mar 26;221(6):jeb173948.

41. Ruhr IM, Takei Y, Grosell M. The role of the rectum in osmoregulation and the potential effect of renoguanylin on SLC26a6 transport activity in the Gulf toadfish (Opsanus beta). Am J Physiol-Regul Integr Comp Physiol. 2016 Jul;311(1):R179–91.

42. Tresguerres M, Levin LR, Buck J, Grosell M. Modulation of NaCl absorption by [HCO3−] in the marine teleost intestine is mediated by soluble adenylyl cyclase. Am J Physiol-Regul Integr Comp Physiol. 2010 Jul;299(1):R62–71.

43. Knobloch S, Skirnisdóttir S, Dubois M, Mayolle L, Kolypczuk L, Leroi F, et al. The gut microbiome of farmed Arctic char (Salvelinus alpinus) is shaped by feeding stage and nutrient presence. FEMS Microbes [Internet]. 2024 Jan 10 [cited 2025 Mar 31];5. Available from: 10.1093/femsmc/xtae011

44. Li Y, Bruni L, Jaramillo-Torres A, Gajardo K, Kortner TM, Krogdahl Å. Differential response of digesta- and mucosa-associated intestinal microbiota to dietary insect meal during the seawater phase of Atlantic salmon. Anim Microbiome. 2021 Jan 7;3(1):8.

45. de Jesús Chavarín-Meza A, Gómez-Gil B, González-Castillo A. Phylogenomic analysis of the Ponticus clade: strains isolated from the spotted rose snapper (Lutjanus guttatus). Antonie Van Leeuwenhoek. 2024 Mar 20;117(1):59.

46. Hickey ME, Lee JL. A comprehensive review of Vibrio (Listonella) anguillarum: ecology, pathology and prevention. Rev Aquac. 2018;10(3):585–610.

47. Rivas AJ, Lemos ML, Osorio CR. Photobacterium damselae subsp. damselae, a bacterium pathogenic for marine animals and humans. Front Microbiol [Internet]. 2013 Sep 25 [cited 2025 Mar 31];4. Available from: https://www.frontiersin.org/journals/microbiology/articles/10.3389/fmicb.2013.00283/full

48. Serracca L, Ercolini C, Rossini I, Battistini R, Giorgi I, Prearo M. Occurrence of both subspecies of Photobacterium damselae in mullets collected in the river Magra (Italy). Can J Microbiol. 2011 May;57(5):437–40.

49. Sheu SY, Jiang SR, Chen CA, Wang JT, Chen WM. Vibrio stylophorae sp. nov., isolated from the reef-building coral Stylophora pistillata. Int J Syst Evol Microbiol. 2011;61(9):2180–5.

50. Mobley HLT. Urease. In: Helicobacter pylori [Internet]. John Wiley & Sons, Ltd; 2001 [cited 2025 Mar 31]. p. 177–91. Available from: https://onlinelibrary.wiley.com/doi/abs/10.1128/9781555818005.ch16

51. Magariños B, Toranzo AE, Romalde JL. Phenotypic and pathobiological characteristics of *Pasteurella piscicida*. Annu Rev Fish Dis. 1996 Jan 1;6:41–64.

52. Rivadeneyra MA, Delgado R, del Moral A, Ferrer MR, Ramos-Cormenzana A. Precipatation of calcium carbonate by Vibrio spp. from an inland saltern. FEMS Microbiol Ecol. 1994 Jan 1;13(3):197–204.

53. Apprill A, Mcnally S, Parsons R, Weber L. Minor revision to V4 region SSU rRNA 806R gene primer greatly increases detection of SAR11 bacterioplankton. Aquat Microb Ecol. 2015;75(2):129–37.

54. Parada AE, Needham DM, Fuhrman JA. Every base matters: assessing small subunit rRNA primers for marine microbiomes with mock communities, time series and global field samples. Environ Microbiol. 2016 May;18(5):1403–14.

55. Martin M. Cutadapt removes adapter sequences from high-throughput sequencing reads. EMBnet.journal. 2011 May 2;17(1):10.

56. Callahan BJ, Mcmurdie PJ, Rosen MJ, Han AW, A AJ. DADA2: High resolution sample inference from Illumina amplicon data. Nat Methods. 2016;13(7):581–3.

57. Yilmaz P, Parfrey LW, Yarza P, Gerken J, Pruesse E, Quast C, et al. The SILVA and “All-species Living Tree Project (LTP)” taxonomic frameworks. Nucleic Acids Res. 2014 Jan;42(D1):D643–8.

58. McMurdie PJ, Holmes S. Phyloseq: An R Package for Reproducible Interactive Analysis and Graphics of Microbiome Census Data. PLoS ONE. 2013;8(4).

59. Wickham H, Averick M, Bryan J, Chang W, McGowan L, François R, et al. Welcome to the Tidyverse. J Open Source Softw. 2019;4(43):1686.

60. Oksanen J, Blanchet F. Package “vegan.” cran.ism.ac.jp. 2022;

61. Andersen K, Kirkegaard R, Karst S, Albertsen M. ampvis2: an R package to analyse and visualise 16S rRNA amplicon data. bioRxiv. 2018 Apr 11;299537.

62. Lin H, Peddada SD. Analysis of compositions of microbiomes with bias correction. Nat Commun. 2020 Jul 14;11(1):3514.

63. Douglas GM, Maffei VJ, Zaneveld JR, Yurgel SN, Brown JR, Taylor CM, et al. PICRUSt2 for prediction of metagenome functions. Nat Biotechnol. 2020 Jun;38(6):685–8.

64. Katoh K, Standley DM. MAFFT multiple sequence alignment software version 7: Improvements in performance and usability. Mol Biol Evol. 2013;30(4):772–80.

65. Larsson A. AliView: a fast and lightweight alignment viewer and editor for large datasets. Bioinformatics. 2014 Nov 15;30(22):3276–8.

66. Capella-Gutiérrez S, Silla-Martínez JM, Gabaldón T. trimAl: A tool for automated alignment trimming in large-scale phylogenetic analyses. Bioinformatics. 2009;25(15):1972–3.

67. Stamatakis A. RAxML version 8: A tool for phylogenetic analysis and post-analysis of large phylogenies. Bioinformatics. 2014;30(9):1312–3.

68. F K. Trim Galore!: A wrapper around Cutadapt and FastQC to consistently apply adapter and quality trimming to FastQ files, with extra functionality for RRBS data. Babraham Inst [Internet]. 2015 [cited 2024 Feb 20]; Available from: https://cir.nii.ac.jp/crid/1370294643762929691

69. Kopylova E, Noé L, Touzet H. SortMeRNA: fast and accurate filtering of ribosomal RNAs in metatranscriptomic data. Bioinformatics. 2012 Dec 1;28(24):3211–7.

70. Kron NS, Young BD, Drown MK, McDonald MD. Long-read de novo genome assembly of Gulf toadfish (Opsanus beta). BMC Genomics. 2024 Sep 18;25(1):871.

71. Dobin A, Davis CA, Schlesinger F, Drenkow J, Zaleski C, Jha S, et al. STAR: Ultrafast universal RNA-seq aligner. Bioinformatics. 2013;29(1):15–21.

72. Bushmanova E, Antipov D, Lapidus A, Prjibelski AD. rnaSPAdes: a de novo transcriptome assembler and its application to RNA-Seq data. GigaScience. 2019 Sep 1;8(9):giz100.

73. Home · TransDecoder/TransDecoder Wiki [Internet]. [cited 2023 Sep 13]. Available from: https://github.com/TransDecoder/TransDecoder/wiki

74. Camacho C, Coulouris G, Avagyan V, Ma N, Papadopoulos J, Bealer K, et al. BLAST+: architecture and applications. BMC Bioinformatics. 2009 Dec 15;10(1):421.

75. Zhang Z, Wood WI. A profile hidden Markov model for signal peptides generated by HMMER. Bioinformatics. 2003;19(2):307–8.

76. Cantalapiedra CP, Hern A, Huerta-Cepas J. eggNOG-mapper v2: Functional Annotation, Orthology Assignments, and Domain Prediction at the Metagenomic Scale. :5.

77. Price MN, Dehal PS, Arkin AP. FastTree: Computing Large Minimum Evolution Trees with Profiles instead of a Distance Matrix. Mol Biol Evol. 2009 Jul 1;26(7):1641–50.

